# Interrogating Mutant Allele Expression via Customized Reference Genomes to Define Influential Cancer Mutations

**DOI:** 10.1101/394627

**Authors:** Adam Grant, Paris Vail, Megha Padi, Agnieszka K. Witkiewicz, Erik S. Knudsen

## Abstract

Genetic alterations are essential for cancer initiation and progression. However, differentiating mutations that drive the tumor phenotype from mutations that do not affect tumor fitness remains a fundamental challenge in cancer biology. To better understand the impact of a given mutation within cancer, RNA-sequencing data was used to categorize mutations based on their allelic expression. For this purpose, we developed the MAXX (**M**utation **A**llelic E**x**pression E**x**tractor) software, which is highly effective at delineating the allelic expression of both single nucleotide variants and small insertions and deletions. Results from MAXX demonstrated that mutations can be separated into three groups based on their expression of the mutant allele, lack of expression from both alleles, or expression of only the wild-type allele. By taking into consideration the allelic expression patterns of genes that are mutated in PDAC, it was possible to increase the sensitivity of widely used driver mutation detection methods, as well as identify subtypes that have prognostic significance and are associated with sensitivity to select classes of therapeutic agents in cell culture. Thus, differentiating mutations based on their mutant allele expression via MAXX represents a means to parse somatic variants in tumor genomes, helping to elucidate of a gene’s respective role in cancer.

## Introduction

Cancer is a complex disease, initiated by DNA alterations within genes that control multiple hallmarks of tumorigenesis, such as deregulated cell growth and genomic instability ^1-3^. Once the genomic architecture of a cancer cell is established, the cancer will continue to evolve to overcome additional regulatory mechanisms and eventually acquire the ability to progress to metastatic disease^4,5^. A primary goal in cancer therapeutics is to target selective pathways that are critical to the tumor’s growth and sustainability^6,7^. However, during tumorigenesis, not only do mutations responsible for the cancer phenotype arise, but so do random mutations that have no effect on the fitness of the tumor. The difficulty in distinguishing driver and passenger mutations represents a core challenge in distilling meaningful functional information from tumor sequencing data^8,9^.

Many efforts are ongoing to determine which mutations contribute to the initiation and progression of cancer. To consolidate information regarding a mutation’s impact on the progression of cancer, well curated databases such as COSMIC^10^ have been established. However, these databases are incomplete and depend heavily on a combination of DNA sequencing and computational software to identify a mutation’s potential contribution towards the development of a tumor^11^. Most driver detection software is centered around one or a combination of three approaches: identifying genes that have an increased mutation rate among a subset of tumor samples^12^, evaluating the functional impact of the mutation^13-15^, and using network analysis to identify gene interactions that have increased mutation rates^16,17^. These computational approaches have contributed a great deal to our current understanding of how and which mutations are involved in the progression of cancer. However, there is a lack of concordance between positive results among the driver mutation identification approaches^18^. Also, a recent evaluation of commonly used driver mutation software demonstrated that in most cases, these approaches have high false positive discovery rates^19^.

One major limitation of established driver identification software is their principle focus on data derived from DNA sequencing and inferred presence of the protein. These approaches provide insight on the distribution of the mutation in cancer and potential function but do not incorporate crucial information on the transcription of the mutated gene. Assumptions on the presence of mutated transcripts can cause inaccurate conclusions as to the mutation’s impact within the tumor. Recent studies have demonstrated that integration of exome and transcriptome genomic profiles can reinforce the interpretation of mutation events ^20,21^. Thus, to increase the understanding of somatic mutations in cancer, we combined exome and RNA-sequencing data to segregate tumor mutations based on their allelic expression.

To enhance mutation allelic expression detection, the software MAXX (**M**utation **A**llelic E**x**pression E**x**tractor) was developed. MAXX has the capability to identify the allelic expression for small insertion and deletion (indel) mutations and single nucleotide variants (SNV), while maintaining high alignment precision. Accurate alignment of RNA-seq reads sets up the groundwork to detect correct RNA allele frequencies for genetic variants^22,23^. Evaluation of the allelic expression for thousands of mutations derived from pancreatic adenocarcinoma (PDAC) established that mutations can be separated into three expression groups based on the presence of RNA-sequencing reads from the mutant allele, wild type allele, or both alleles. These three mutation expression groups were established to have unique biological features that revealed expected characteristics of driver mutations, passenger mutations, and a less appreciated group of mutations that appeared to be selectively silenced within the tumor context. Mutation expression groups also assisted in the identification of PDAC subtypes that have prognostic relevance.

## Results

### Developing an unbiased tool to assess the allelic expression of somatic variants

While there are multiple approaches to delineating if a particular mutation is expressed, much of this work has involved gene-specific analysis instead of a holistic evaluation of the cancer genome. To attain a global assessment of mutation expression within tumors, we quantified the allelic expression of thousands of mutations (missense, nonsense, and small insertion/deletion) derived from PDAC patient-derived cell lines, patient-derived xenografts (PDX), and primary tumors obtained from The Cancer Genome Atlas (TCGA)^24,25^. Because mutations can exist as either somatic single nucleotide variant (SNV) or small insertion/deletion (indel), a unique approach was developed to confidently identify the allelic expression of all mutations. Rather than depend on tumor RNA-sequencing reads to be correctly aligned onto a standardized reference genome (e.g. Hg19), the MAXX (MAXX: **M**utation **A**llelic E**x**pression E**x**tractor) method creates a new reference genome that is tumor selective and specifically extracts mutant allele expression. The MAXX approach utilizes mutation calls derived from DNA sequencing to generate a tumor-specific reference genome (Fig. 1a). This precise reference genome was then used in conjunction with Tophat2^26^ to enhance the alignment of the tumor’s RNA-sequencing reads to the mutant allele. The approach of MAXX enables the calculation of precise RNA mutant allele frequencies for essentially any variant type that is defined by the DNA sequencing approach, allowing a comprehensive and unbiased analysis of mutation allelic expression.

**Fig. 1:**
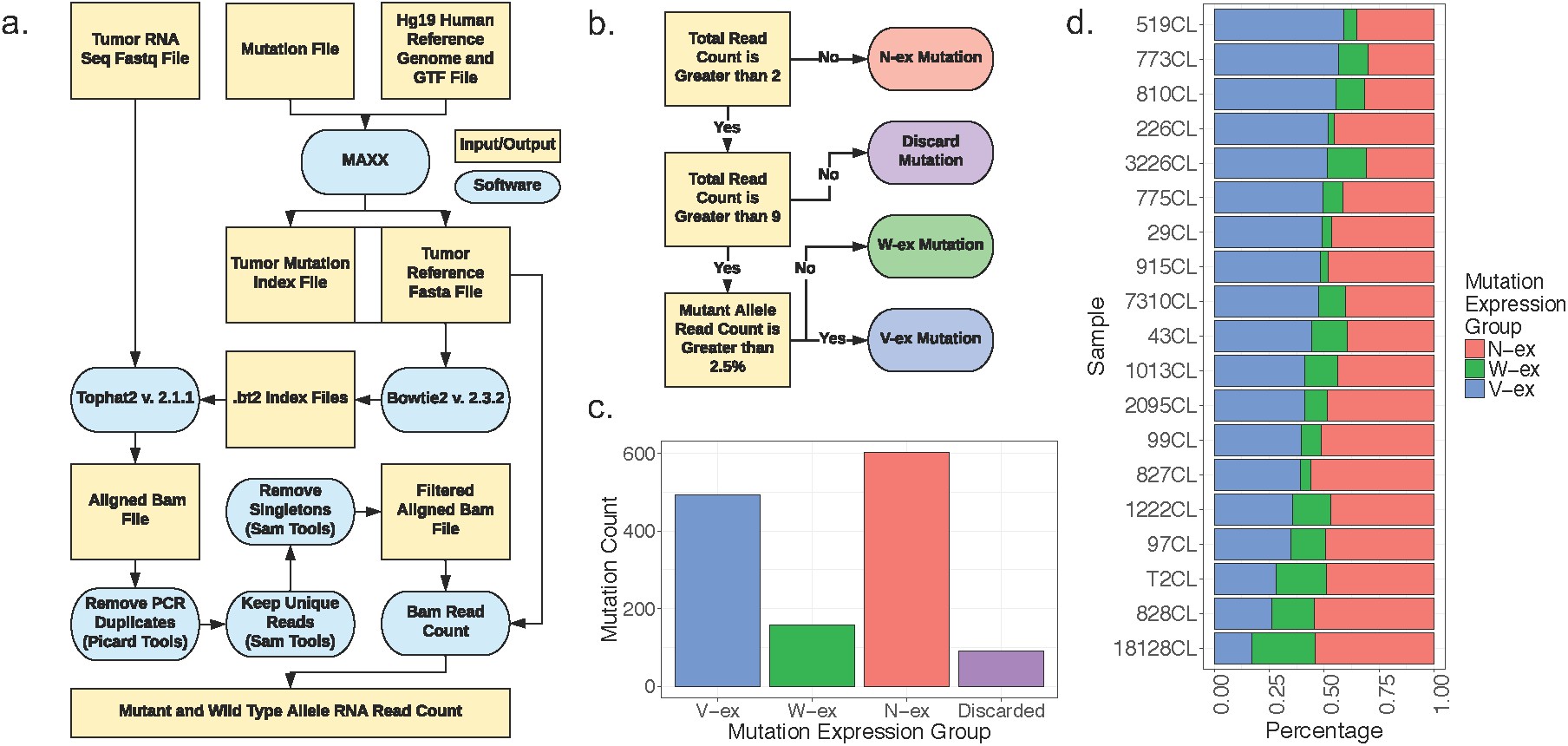
Developing an unbiased tool to assess the expression of somatic variants. (a) Flowchart of the MAXX pipeline, which ultimately identifies the RNA read count for the mutant allele and the wild type allele. (b) Methodology for mutation expression group placement, represented as V-ex (blue), W-ex (green), and N-ex (red): V-ex, mutations that express the mutant allele; W-ex, mutations that only express the wild type allele; N-ex, mutations that don’t express the wild type allele or the mutant allele. (c) The number of mutations associated with each mutation expression group. These mutations were derived from 19 PDAC patient derived cell lines. (d) Individual distributions of mutation expression groups for each patient derived cell line.

### Different patterns of variant allelic expression in tumor models

To initially interrogate the efficacy of the MAXX pipeline, we generated mutation expression profiles, a file containing the DNA and RNA allele frequencies for all identified mutations, for 19 PDAC patient derived cell lines. These cell lines underwent both RNA-sequencing and exome sequencing relative to the normal tissue. Cell line models were ideal for the primary evaluation of MAXX because of the absence of stromal contamination, substantially decreasing the confounding feature of tumor purity^27,28^. Assessment of the 19 mutation expression profiles identified that tumor mutations can be classified based on the number of reads that align to the wild type allele, mutant allele, or both alleles (Fig. 1b). In order to prevent misclassification between mutations that do and do not express the mutant allele, mutations that had a total RNA read count between 3 and 9 were discarded from the study. All non-discarded mutations were placed into one of the following three mutation expression groups: 1. Mutations that express the variant allele, labeled as the variant expressed group (V-ex, Blue) 2. Mutations wherein only the wild type allele is expressed, labeled as the wild type expressed group (W-ex, Green). 3. Mutations that do not express the wild type or the mutant allele, labeled as the not expressed group (N-ex, Red). These three mutation expression groups have also been identified in patient derived cell lines from myeloma patients^29^. The distribution of the three mutation expression groups was summarized across all patient derived cell lines (Fig. 1c). The N-ex group made up half of all the variants, while only a small subset of mutations fell into the W-ex group. This overall distribution was observed in all individual models, suggesting that the expression distribution is not specific to a given sequencing run or tumor (Fig. 1d).

### MAXX pipeline accurately maps indels and is computationally efficient

Previous studies have shown that RNA-sequencing aligners are capable of aligning SNV, but have poor alignment precision for reads containing an indel mutation^30,31^. However, the unique workflow of MAXX allows the allelic expression to be confidently calculated for both SNV and indel mutations (Fig. 2a). To enhance indel alignment for RNA-sequencing data, which is a critical aspect in identifying accurate RNA allele frequencies^20^, MAXX generates a tumor-specific reference genome based on mutation calls derived from DNA-sequencing. When comparing RNA allele frequencies derived from bam files that were aligned with Tophat2 using either a MAXX or Hg19 reference genome and underwent important filtering steps for allelic expression analysis^21,32^, it was identified that the MAXX reference genome significantly enhanced the allelic expression of indel mutations, compared to the Hg19 reference genome (Fig. 2b). There was relatively little variation between SNV RNA allele frequencies using the MAXX or Hg19 reference genomes.

**Fig. 2:**
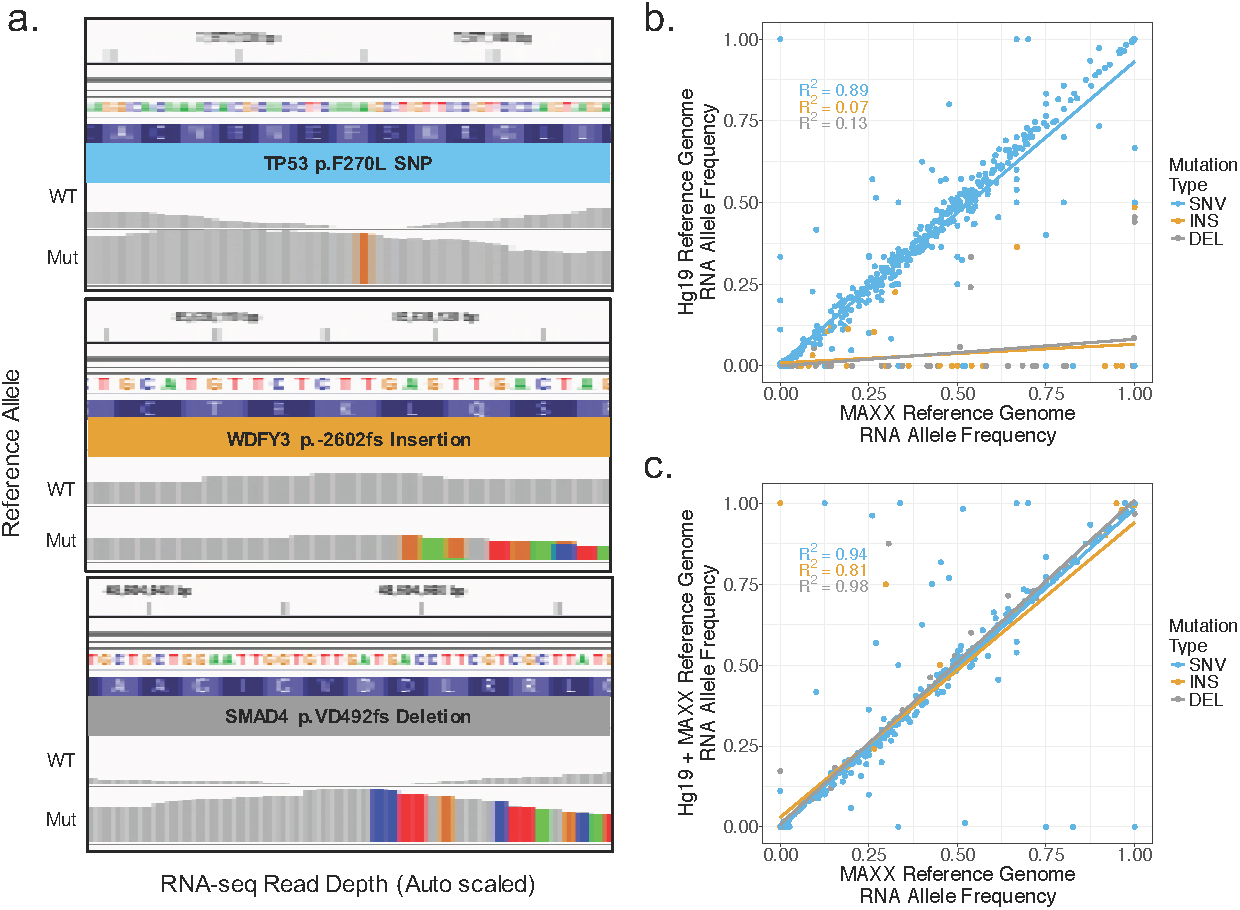
MAXX pipeline accurately maps indels and is computationally efficient. (a) Integrative genomic viewer visualization of raw RNA-seq reads that aligned to the wild type allele or the mutant allele for a SNV, insertion, and deletion mutation. Non-gray colors represent an alternative nucleotide compared to the reference genome (b) Comparison of the RNA mutant allele frequencies calculated by using either the MAXX generated reference genome or the Hg19 reference genome. (c) The contrast between the RNA mutant allele frequencies identified by using either the MAXX generated reference genome or the Hg19 reference genome with an appended mutant genome created by MAXX.

Generating a reference genome for each tumor sample would require a significant amount of computational processing and storage for large studies. However, MAXX generate reference genomes are significantly smaller than the Hg19 reference genome. This decrease in size is due to the tumor reference genome containing only the wild type and mutant sequence for each gene harboring a mutation. Using a reference genome containing only the mutated genes significantly decreases the storage space and computational resources used to index the reference genome and run Tophat2 compared to the Hg19 reference genome (Supplementary Fig. 1). To determine if there is over selection of aligned reads due to the highly truncated genome, we compared RNA mutant allele frequencies established from an Hg19 reference genome with the appended MAXX mutant genome to a MAXX generated reference genome (Fig. 2c). These two reference genomes delivered veritably identical RNA mutant allele frequencies and placement of mutations into expression groups (Supplementary Fig. 2). Overall, this demonstrates that a simplified reference genome provides a robust substrate for differential alignment of transcript reads and can confidently be used to generate mutation expression profiles.

### Mutant allele expression is associated with the DNA mutant allele frequency but not transcript expression level

To resolve if the RNA mutant allele frequency is associated with the DNA mutant allele frequency in PDAC, we evaluated the dual relationship across the three mutation expression groups (Fig. 3a.) In general, the variant allele expression of V-ex mutations correlated with their DNA allele frequency (R^2^=.59). This correlation was similarly identified in another study using CT26, B16F10, and 4T1 mouse cell lines^33^. However, in our dataset it was observed that multiple genes with a V-ex mutation had a 100% RNA mutant allele frequency, despite a DNA mutant allele frequency of approximately 50%. This discrepancy between DNA and RNA mutant allele frequencies suggests that the non-mutated allele of these genes is selectively silenced, which is a common phenomenon for tumor suppressors. N-ex and W-ex mutation expression groups have no expression of the mutant allele, and as expected they had a RNA mutant allele frequency of zero (Fig. 3b). To determine if a mutation expression group was biased towards a specific mutation type (missense, deletion, insertion, nonsense), as might be expected from processes such as nonsense mediated decay, the proportion of mutation types was measured between mutation expression groups (Supplementary Fig. 3). Statistical testing illustrated that there is no correlation between the mutation type and mutation expression groups. Interestingly, the variant allele frequency of mutation expression groups established that mutation expression provides insight on mutation selectivity (Fig. 3c). The V-ex group has a significantly higher mean variant allele frequency than the other two groups, suggesting that clonal drivers of the tumor are generally expressed. As for the N-ex group, most mutations had a variant allele frequency between 30% and 40%, suggesting that many of these mutations, whilst not expressed, are in fact largely clonal in the tumor. In contrast, the mean variant allele frequency of the W-ex group is considerably lower than the other two mutation expression groups. To interrogate if there is some selective feature of W-ex mutations that leads to message loss, we compared the mean gene expression of samples that did not have a mutated allele to the mean gene expression of samples that did contain a mutated allele (Fig. 3d). To obtain the global gene expression for all 19 patient derived cell lines, raw read counts were obtained using HTSeq^34^ on BAM files aligned to the Hg19 reference, then normalized using edgeR^35^. The overall expression of genes that contained a N-ex mutation is substantially less than genes with a V-ex or W-ex mutation, as expected. This confirms that genes with a N-ex mutation are rarely expressed, and thus mutations within this class likely have little influence on PDAC. On the contrary, genes with a V-ex or W-ex mutation contained a relatively high gene expression within the tumor, suggesting that genes with a V-ex or W-ex mutation are important to the progression of the tumor. A two-tail paired t-test between the mean gene expression of mutated and non-mutated genes was performed for each mutant expression group (Fig. 3e). While significant differences were observed between mutated and unmutated genes among each expression group, the magnitude change was marginal. To determine if an increased gene expression, yet lack of mutant allele expression of W-ex mutations was due to RNA splicing events, the exon expression of mutated and non-mutated exons was quantified using DEXseq^36^ and normalized using edgeR, then compared in a similar manner as the transcript data (Supplement Fig. 4). Equivalent to the transcript analysis, there was little variance between the mutated and non-mutated exon expression. These findings imply that there is not a strong selection against transcript or exon expression of the W-ex group.

**Fig. 3:**
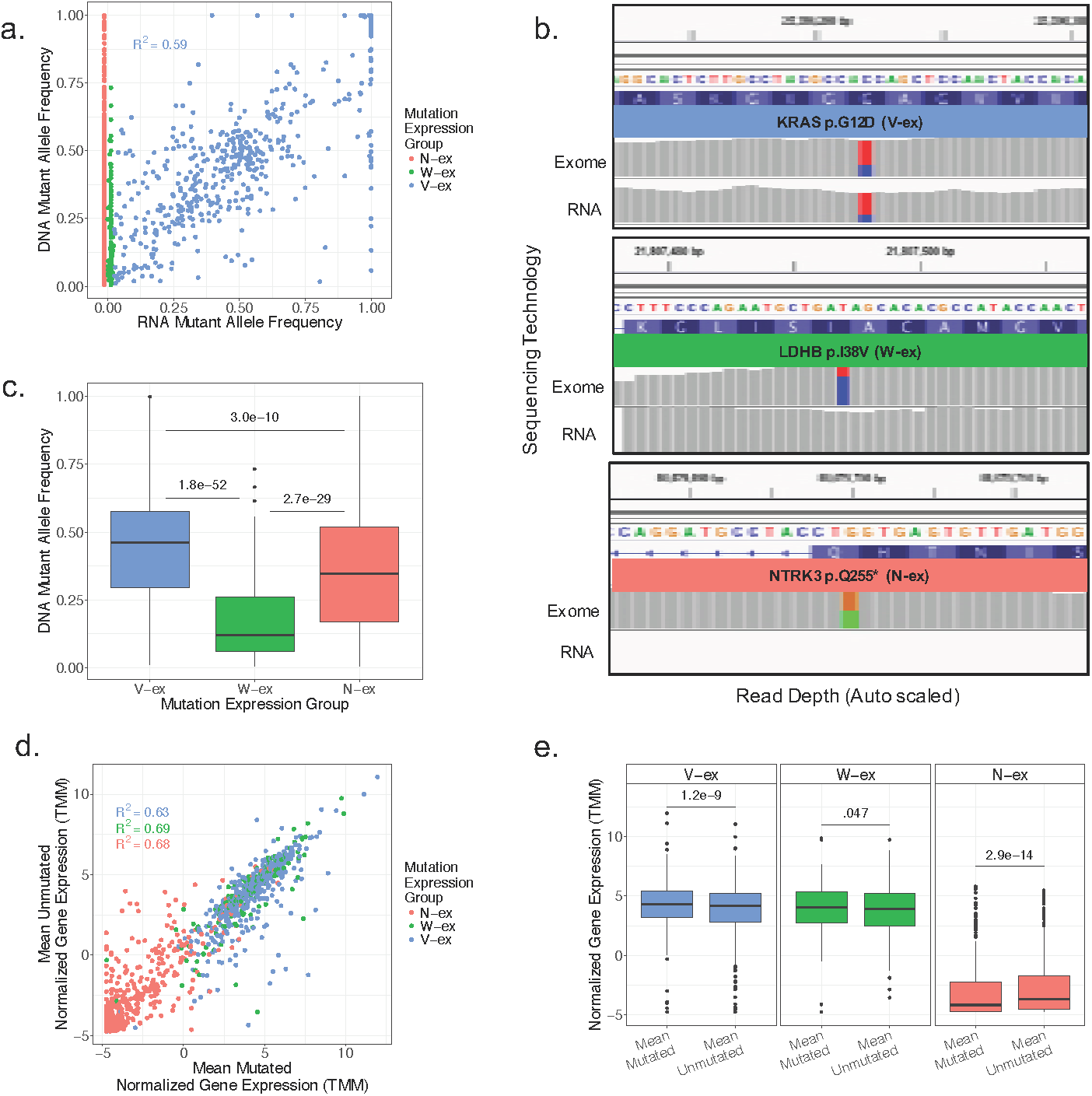
Mutant allele expression is associated with the DNA mutant allele frequency but not transcript expression level. (a) Each mutation’s DNA allele frequency plotted against its corresponding RNA allele frequency. (b) Integrative genomic viewer representation of the exome sequencing and RNA sequencing for a V-ex mutation, W-ex mutation, and N-ex mutation. Non-gray colors represent the presence of conflicting nucleotides aligned to the reference genome. (c) The distribution of DNA allele frequencies for the three mutation expression groups and statistical significance based on a two-sample t-test with a two-tail p-value. (d) Contrast of the mean gene expression levels of samples that do contain the mutated gene to the mean gene expression levels of samples that don’t contain the mutated gene. (e) Two sample paired t-test with a two-tail p-value was performed between of the average gene expression levels of samples with the mutated gene and samples without the mutated gene for each mutation expression group.

### PDAC associated genes mainly fall into the V-ex and W-ex groups

To more fully understand the significance of mutation expression profiles, we evaluated the mutated genes within the three expression groups. In the context of PDAC tumors, it is well known that there are four genes that are frequently mutated and considered disease drivers: KRAS, TP53, SMAD4, and CDKN2A^37^. These four genes were repeatedly observed to contain V-ex mutations, skewing the mutation frequency distribution of the V-ex group as shown in the violin plots (Fig. 4a). In comparison, the frequency distribution of mutated genes in W-ex and N-ex groups is heavily centered around one, implying little recurrence of such genes within PDAC. Oncogenes such as KRAS generally have a DNA mutant allele frequency of approximately 50%, because only one allele is required to be mutated for the gene to act as an oncogene^38^. However, in select cases there is specific selection for the expression of only the mutant allele. Interestingly, this largely occurs due to genetic as opposed to allele specific gene expression (Fig. 4b). In contrast, tumor suppressors TP53, SMAD4, and CDKN2A have mutation allele frequencies that approach 100% and mainly express only the mutated allele. In the case of SMAD4 and CDKN2A, this would be expected. However, as TP53 mutations can have gain of function mutations, this data suggests there is exceedingly strong pressure to lose the wild-type allele during tumor development within the pancreas.

**Fig. 4:**
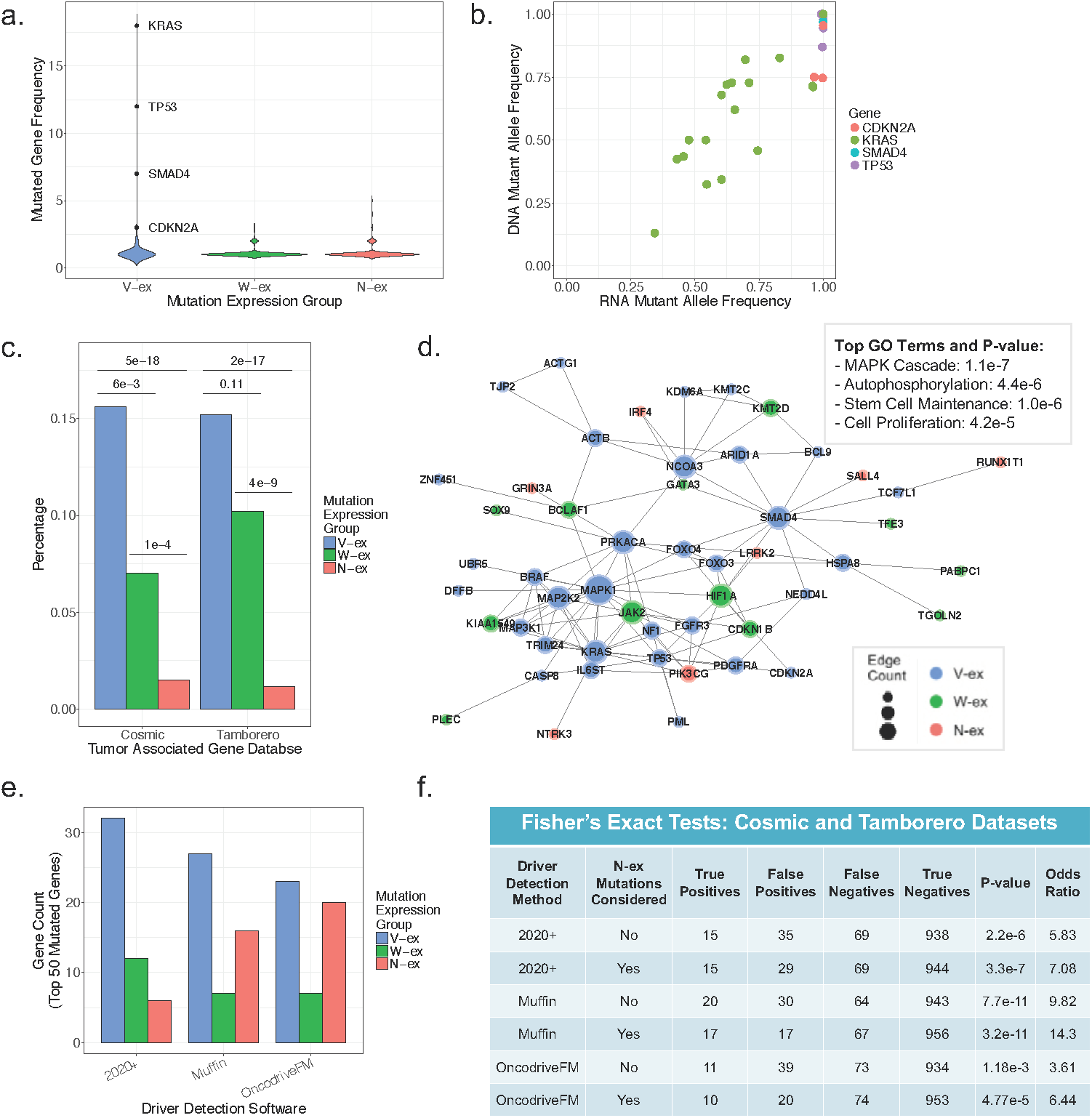
PDAC associated genes mainly fall into the V-ex and W-ex groups. (a) The gene mutation frequency of all V-ex, W-ex and N-ex mutations. (b) Comparison of the DNA allele frequency and RNA allele frequency for the most well-known PDAC mutations (KRAS, CDKN2A, SMAD4, TP53). (c) Percentage of mutations that are associated with the Cosmic and Tamborero cancer gene datasets for each mutation expression group. Statistical significance was performed using a two-proportion z-test between each of the mutation expression groups. (d) A network generated from the V-ex, W-ex, and N-ex mutated genes that were present in either the Cosmic or Tamborero datasets. (e) The distribution of mutation expression groups from the top 50 ranked mutated genes outputted by the driver detection methods 2020+, Muffin, and OncodriveFM. (f) The specificity of the three driver detection methods to identify cancer associated genes from the Cosmic and Tamborero from their 50 top ranked mutated genes. Cancer gene specificity was calculated for either all top 50 mutated genes or the top 50 mutated genes that are not classified as N-ex.

In order to delineate the putative significance to cancer, we concentrated on the percentage of mutations in each mutation expression group that were present in either the COSMIC.v80^10^ or Tamborero *et al.*^39^ cancer associated gene datasets (Fig. 4c). A two-proportion z-test was used to calculate if the proportion of cancer-associated genes was significant between each of the mutation expression groups. As expected, due to the frequency of KRAS, TP53, SMAD4, and CDKN2A, the V-ex group contained the most statistically significant percentage of mutated genes which are associated with cancer. Comparing the W-ex group to the N-ex group demonstrated that the W-ex group had a significantly higher percentage of mutated genes that are associated with cancer. To determine if there is a relationship between mutated cancer associated genes and mutation expression groups, network analysis was performed using the Cytoscape^40^ ReactomeFI plugin^41^ (Fig. 4d). Evaluation of the generated network indicates that cancer associated genes with a W-ex mutation are well integrated with cancer associated genes containing a V-ex mutation. As for genes containing a N-ex mutation, they were commonly found on the outsides of the network, having little contribution to the structure of the network. These results support the hypothesis that W-ex mutations occur in genes that are important to the tumor, despite their lack of clonal selection and mutant allele expression. In contrast, N-ex mutations appear to be in genes that have little to no impact on tumor progression.

Most, if not all, current driver detection methods do not integrate allelic expression information when predicting influential cancer mutations. Thus, to determine if incorporation of mutation allelic expression can increase cancer gene specificity, we started by employing three commonly used driver detection methods. The three methods selected were 2020+^19^, Muffin^17^, and OncodriveFM^15^. These methods were chosen because of their capability to handle indel mutations and relatively small sample sizes. When comparing the mutation expression group distribution of the top 50 ranked mutated genes identified by 2020+, Muffin and OncodriveFM, there was a consistent tradeoff between V-ex and N-ex groups (Fig 4e). The driver detection method 2020+ had the highest proportion of V-ex mutated genes, while OncodriveFM had the lowest proportion of V-ex mutated genes. To test the specificity of each method to identify cancer associated genes from the Cosmic and Tamborero datasets, a Fisher’s Exact test was performed with the assumption that only the top 50 mutated genes predicted by each method were cancer associated (Fig. 4f). Overall, Muffin outperformed 2020+ and OncodriveFM. When we removed the N-ex mutations from the top 50 putative driver mutations, all three methods had an increased precision in predicting cancer associated mutated genes. This suggests that incorporation of mutations’ allelic expression can assist in reducing false positive discovery rates of driver mutation detection methods and significantly improve downstream analyses.

### Conservation of mutational expression features from cell lines to PDX tumors

To determine the efficacy of the MAXX pipeline in producing mutation expression profiles for tumors, the MAXX pipeline was evaluated on PDX models. 8 of the 19 patient derived cell lines have matched PDX models^24^; therefore, these 8 PDX models shared the same tumor specific reference genome as their corresponding cell line. However, these 8 PDX models had their own transcriptome sequenced. Similar to the cell line data, the three mutation expression groups were determined based on wild type and mutant allele read counts, and the RNA mutant allele frequency was plotted against the DNA mutant allele frequency (Fig. 5a). The PDX models demonstrated that mutation expression groups are present in both in-vitro and in-vivo. Equivalent to the cell lines, RNA mutant allele frequency is strongly correlated with DNA mutant allele frequency. Amongst the V-ex mutations, there was exceedingly high commonality in expression features between PDX and cell lines (Fig. 5b). This conservation was even more extreme for N-ex mutations (Fig. 5c). However, W-ex mutations demonstrated more variability between PDXs and their corresponding cell lines, where most of the unique W-ex mutations fell within the PDX samples (Fig. 5d). While evaluating how mutation allele frequency associates with the common vs unique between PDX and cell lines via a two-tailed Wilcoxon Mann Whitney test, V-ex variants demonstrated that there is clearly a relationship (Fig. 5e). Although not statistically significant, W-ex mutations demonstrated an opposite relationship. This result suggests that in part, the lack of conservation of mutation expression is a reflection of the sub-clonal feature of the specific variant.

**Fig. 5:**
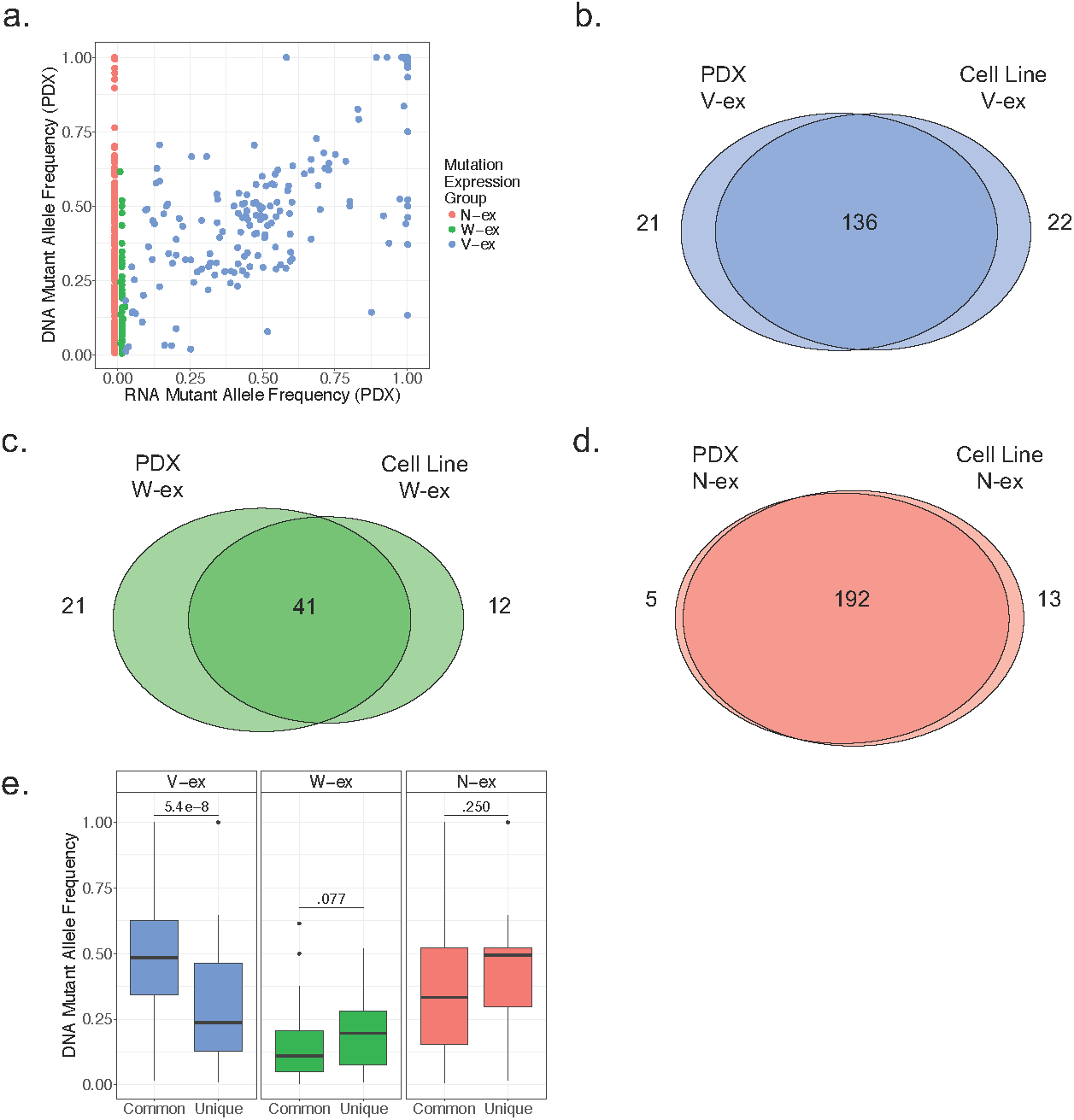
Conservation of mutational expression features from cell lines to PDX tumors. (a) Comparison of the DNA allele frequency and RNA allele frequency of each mutation derived from the PDX samples. (b-d) The overlap of V-ex, W-ex, and N-ex mutations between the corresponding cell line and PDX samples. Mutations classified as “discarded” in either the cell line or PDX data were excluded. (e) A two-tailed Wilcoxon Mann Whitney test between the DNA allele frequency of mutations that are present in both the cell line and PDX samples and the DNA allele frequency of mutations that are unique to either the cell line or PDX samples was performed for each mutation expression group.

### Mutation expression profiles from primary tumors

Primary tumors frequently contain stromal contamination, which decreases the sensitivity of mutation calls and diminishes the RNA-sequencing reads associated with the tumor^42^. Consequently, stromal contamination has been reported to cause inaccurate results when performing DNA methylation and subtyping analyses^28,43^. Thus, to limit the issue of tumor purity, we focused on the 75 TCGA PDAC cases that had the highest tumor purity score determined by the software ESTIMATE^44^ and had a V-ex mutation percentage of at least 10%. Similar to the cell line and PDX models, the three mutation expression groups were present in the TCGA samples (Fig. 6a). However, the distribution of the tumor DNA mutant allele frequency is mostly between 0-50% rather than 0-100% as it was for the cell lines (Fig. 6a vs. 3a). To determine if stromal contamination also altered the RNA allele frequency, we focused on KRAS, CDKN2A, TP53, and SMAD4 mutations (Fig. 6b). While the range of the RNA allele frequency for KRAS and CDKN2A is comparable to that observed in cell line data, the range significantly changed for TP53 and SMAD4. This suggests that SMAD4 and TP53 expression is observed in both the tumor specimen and the stromal compartment. In spite of this complexity, most mutations in these canonical PDAC mutations were identified to be expressed within the tumor, as seen in the cell and PDX data.

**Fig. 6:**
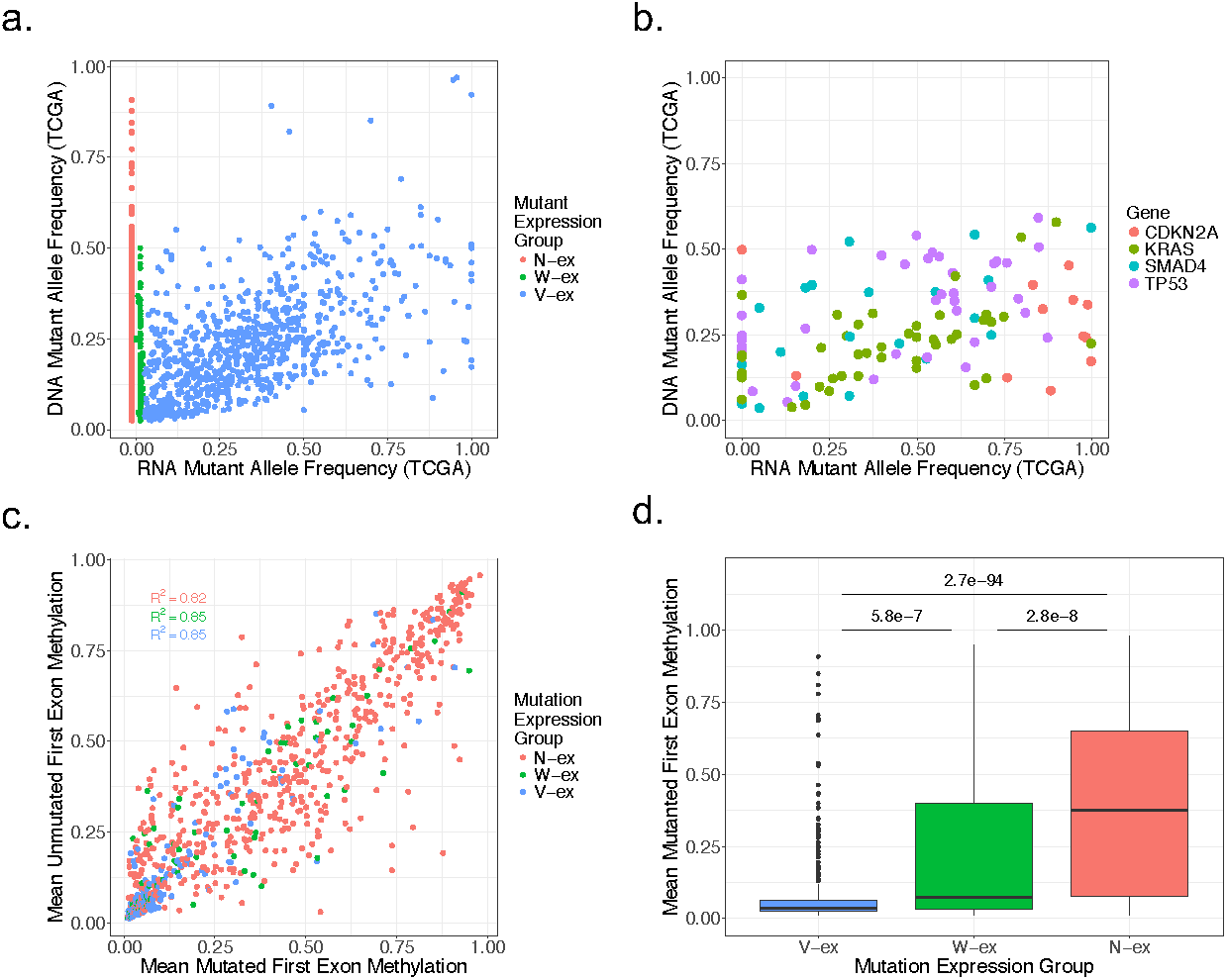
Mutation expression profiles from primary tumors. (a) Comparison of the DNA allele frequency and RNA allele frequency of each mutation derived from the TCGA samples. (b) Correlation of the DNA allele frequency and the RNA allele frequency of CDKN2A, KRAS, SMAD4, and TP53 mutations identified within the TCGA samples. (c) The average first exon methylation of samples that didn’t contain the mutated gene was plotted against the average first exon methylation of samples that did contain the mutated gene. This was performed for non-discarded mutated genes that available first exon methylation data. (d) A two-sample t-test with a two-tail p-value was performed on the mutant first exon methylation between each mutation expression group.

Using mutation expression and methylation profiles from TCGA samples, we were able to interrogate the levels of DNA methylation across mutation expression groups. To measure the methylation of genes containing a mutation, the mean methylation percentage of each recorded CpG within the first exon was calculated^45^. Similar to the mutant gene expression analysis, the mean first exon methylation of samples that did not have a mutation in the gene was compared to the mean first exon methylation of samples that did contain the mutated gene (Fig. 6c). Evaluation of the coefficient of determination (R^2^) for each expression group demonstrated that there is little deviation between wild type and mutated gene first exon methylation. However, when comparing the average mutated first exon methylation between mutation expression groups via a two-tail t-test, the mean first exon methylation of N-ex mutations is significantly higher than the V-ex and W-ex mutations (Fig. 6d). This finding suggests that hypermethylation of the gene is a probable explanation for lack of expression among genes with a N-ex mutation. When comparing the first exon methylation between the V-ex and the W-ex groups, genes containing a W-ex mutation were identified to have a statistically higher first exon methylation. One possible explanation of why the W-ex group had a statistically lower first exon methylation than the N-ex group, yet a statistically higher first exon methylation than the V-ex group is that these genes are susceptible to hemimethylation^46^. Hemimethylation of genes with a W-ex mutation would also explain why the mutant allele transcript from these genes is undetectable.

### Separating mutations by expression significantly enhances PDAC subtyping

In order to determine if mutation expression groups provide insight into prognostic relevance, the Network Based Stratification (NBS)^47^ pipeline was utilized on both the 19 patient-derived cell lines and 75 TCGA samples, as implemented by Morvan et al^48^. NBS uses a predefined protein-protein interaction network to separate samples into a predefined number of subtypes, based on a mutation sample matrix. To quantify the prognostic capabilities of mutation expression groups, a survival log rank statistic-Log10(p-value) was calculated for only the TCGA samples. The log rank statistic p-value was calculated for all possible combinations of mutation expression groups, using 3-8 predefined number of subtypes (Fig. 7a). The combination of mutation expression groups that had the strongest statistical prediction of prognosis was the V-ex and W-ex mutations, and this held true regardless of the number of predefined subtypes. These findings further support that both the V-ex and W-ex mutations are relevant to tumor biology and occur within similar portions of the protein-protein interaction network. Comparatively, N-ex mutations have little influence on the progression of the tumor and appear to be randomly distributed within a protein-protein interaction network. To compare the prognostic prediction of the V-ex and W-ex mutations to gene rankings of popular driver detection methods, we performed a similar NBS analysis on results produced by 2020+, Muffin and OncodriverFM (Fig. 7b). To perform an unbiased comparison between all methods, the same input data was used for each method to produce a ranked list of mutated genes. Then for the NBS analysis, the same number of ranked mutated genes was used to generate a mutation sample matrix (n = 1,349). Compared to each driver detection method, the combination of the V-ex and W-ex mutations provided the best prognostic prediction for each number of predefined subtypes. Thus, these results imply that that similar to Fig. 4f, current driver detection methods report many false positives which contaminate results and affect downstream analyses. However, by separating mutations based on their allelic expression, it was possible to remove a large subset of mutations that appear to have little to none influence on the progression of the tumor and significantly enhance prognosis prediction with an emphasis on mutation data.

**Fig. 7:**
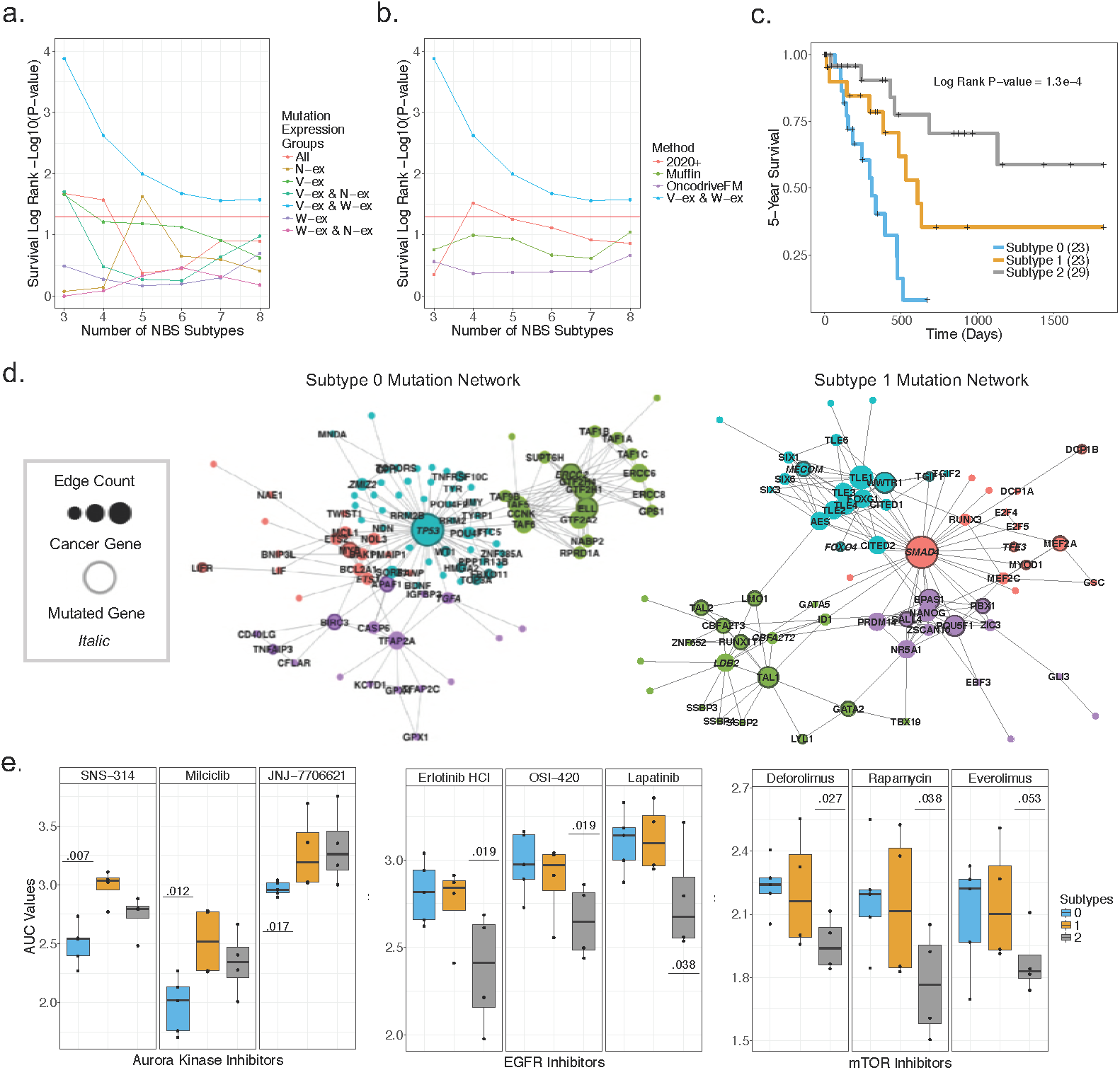
Separating mutations by expression enhances PDAC subtyping. (a) The log rank statistic-log10(p-value) based on the NBS results for 75 TCGA samples. The log rank statistic-log10(p-value) was calculated for the NBS results of all pairwise combination of mutation expression groups from 3-8 number of predefined subtypes. (b) The log rank statistic-log10(p-value) based on the NBS results for 75 TCGA samples using the stratified samples based on results from 2020+, Muffin, OncodriveFM or V-ex and W-ex mutations. (c) Kaplan Meier plot for TCGA samples based on the NBS results using V-ex and W-ex mutations with number of subtypes equal to three. (d) Networks signifying the mutated pathways that are unique to subtype 0 and subtype 1. (e) Cell line drug response data, grouped by identified subtypes and statistical significance via a Wilcoxon Mann Whitney less than test.

The most statistically significant number of stratified subtypes based on the V-ex and W-ex groups was three. The Kaplan-Meier plot generated from this result is shown (Fig. 7c). To determine which genes were unique to the newly established subtypes, a Wilcoxon Mann Whitney greater than test was performed on the NBS gene smooth scores between each two-way comparison of the three subtypes (*e.g.* subtype 0 vs subtype 1 and subtype 2). After adjusting the p-values using the Benjamini/Yekutieli method^49^, networks were generated for each subtype using genes that contained an adjusted p-value less than .01 (Fig. 7d). Subtype 0 and subtype 1 had uniquely defined networks, centered around TP53 and SMAD4 respectively. GO term analysis identified that subtype 0 mutations influence apoptosis pathways, while subtype 1 mutations are involved in cell differentiation pathways. In contrast, samples classified as subtype 2 had only a few statistically significant mutated interactions and thus a well-structured network could not be generated. Interestingly, the sole mutation of either TP53 or SMAD4 was not sufficient to subtype the samples used in this study (Supplementary Fig. 5). This data supports the use of pathway and MAXX analysis to decipher new prognostic subtypes based on mutation data.

In order to evaluate if such subtyping could have relevance to treatment of PDAC, statistical analysis was performed using drug response data from 13 of our 19 patent derived cell lines (Fig. 7e). Among the cell lines with drug sensitivity data, five were categorized as subtype 0, four were categorized as subtype 1, and four were categorized as subtype 2. To determine the significance of drug treatments between subtypes, a Wilcoxon Mann Whitney less than test was performed on the drug AUC values between each two-way comparison of the three subtypes. The drugs that were most consistent between subtype 0 and subtype 2 were, respectively, aurora kinase and EGFR inhibitors. Subtype 2 was also identified to respond better to mTOR inhibitors relative to the other subtypes. Unexpectedly, subtype 1 did not display selective sensitivity to any drug treatments. These results suggest that MAXX-derived data can be applied to predict drug sensitivity, in addition to prognosis.

## Discussion

This study shows that tumor specific reference genomes generated via MAXX provide a substrate for a comprehensive analysis of mutation allelic expression. In comparison to standardized reference genomes, tumor specific reference genomes significantly enhance RNA-sequencing alignment of reads containing indel mutations. This feature allows accurate allelic expression to be detected for all mutation types. In the context of our analysis using 19 patient derived cell lines, the capability to confidently detect allelic expression of indel mutations increased the mutation sample size by approximately 20%. MAXX also provided an unbiased allelic expression analysis among genes that commonly exhibit both SNV and indel variants (e.g. TP53 and SMAD4). The robustness of MAXX was measured on patient derived cell lines, PDX models, and primary tumors. Performing these analyses among different tumor models demonstrated that the MAXX pipeline is a comprehensive, consistent and computationally efficient method to identify mutation allelic expression. MAXX is also capable of generating customizable gene specific reference genomes that can be used as input for an alignment software to effectively query DNA or RNA allele frequencies for specific gene mutations from any organism with a reference genome and GTF file.

Centered on variants derived from PDAC, we report the discovery that mutation allelic expression can be used to separate mutations based on their respective impact on the tumor phenotype. The mutant expression group that was mainly responsible for the progression and clonality of the tumor was the V-ex group. This expression group consisted of not only the major PDAC oncogene (KRAS), but also all of the well-known PDAC tumor suppressor genes (TP53, SMAD4, and CDKN2A). Regardless of their assumed loss of function, our data identified that it is common for tumor suppressors to be transcriptionally present. This finding supports that RNA allele frequencies derived from allelic expression analysis can effectively be used to identify bona fide tumor suppressors. To illustrate, the well-known tumor suppressor gene SMAD4 commonly loses its expression of the wild type allele due to a homozygous deletion. Consequently, SMAD4’s role as a tumor suppressor can be identified using both DNA mutant allele frequencies and RNA mutant allele frequencies. However, within our cell and PDX data, it was observed that multiple mutated genes had a RNA mutant allele frequency of 100% regardless of a DNA mutant allele frequency of 50%. Thus, MAXX derived RNA mutant allele frequencies are more suited to identify genes that lose their expression of the wild type allele, which is an expected feature of tumor suppressor genes, compared to DNA mutant allele frequencies. Two genes from our data set that had an approximate 50% DNA allele frequency, a 100% RNA allele frequency, and have been demonstrated to act as tumor suppressors within the literature were DUSP5^50^ and EI24^51^. Additional genes that had similar DNA and RNA frequency as DUSP5 and EI24 but no previous research on their potential role as a tumor suppressor were UBXN11, CAPN15, C6orf62, GPRIN1, KIAA2018, PCDHGA10, and PCGF1.

The mutation expression group that appeared to have little effect on tumor progression was the N-ex group. Despite their relatively high mutant allele frequency, N-ex mutations were identified to be within genes that had a significantly lower transcript level than genes containing a V-ex or W-ex mutation. Using TCGA methylation data, we identified that a large proportion of genes containing a N-ex mutation had a hypermethylated first exon, regardless of the presence or absence of a mutation. This finding suggests that genes harboring a N-ex mutation are genes that are typically the target of epigenetic silencing across pancreatic cancer. Thus, the minimal impact of N-ex mutations on tumor progression is partially explained by their occurrence within genes that are developmentally silenced or are within genes that are more susceptible to transcription silencing among tumor samples. We observed that some N-ex classified genes have been previously identified to be associated with cancer. However, in comparison to cancer associated genes with a V-ex or W-ex mutation, cancer associated genes with a N-ex mutation had little impact on well-characterized mutated PDAC pathways such as MAPK cascade, autophosphorylation and stem cell maintenance. It was also observed, that when taking into consideration whether a mutation was classified as a N-ex mutation, the mutation detection methods 2020+, Muffin, and OncodriveFM had an increased specificity for pre-classified cancer associated genes. Thus, we suggest that unexpressed genes that are the target of a mutation provide little insight into the current progression and therapeutic response of the tumor, relative to expressed genes that contain a mutation.

In addition to identifying groups of mutations that resemble driver mutations (V-ex) or passenger mutations (N-ex), we found a third mutation group, the W-ex group, which appeared to be an understudied subset of mutations. W-ex mutations were distinguished by their absence of the mutant allele transcript. The inability to detect the transcript of the mutant allele can possibly be due to low DNA mutant allele frequencies, which confounds sufficient RNA-seq reads.However, we are confident that a majority of the W-ex classified mutations identified in the cell line and PDX models are true positives. This is because the patient derived cell lines which were used in both the exome and RNA sequencing originated from the same portion of the tumor. Also, each cell line and PDX sample had their RNA sequenced in triplicate. As for the TCGA data, it is unknown whether the same portions of the tumor were used for both exome and RNA sequencing; thus, sampling could factor into the detection of W-ex mutations. Nonetheless, a number of W-ex classified mutations within the TCGA data had a DNA allele frequency greater than 20% and a substantial depth of RNA-sequencing reads. For example, the mutated gene FN1 had a DNA allele frequency of 50% and 99 reads that aligned to the wild type allele. Thus, it appears that there is a subset of mutations wherein only the wild type allele is expressed.

Despite the lack of the mutant allele transcripts, W-ex mutations appear to be biologically relevant to the progression of the tumor, similar to V-ex mutations. In comparison tothe N-ex group, the W-ex group had a significant proportion of cancer associated genes and high gene expression levels. However, W-ex mutations were more sub clonal and infrequent within tumor samples relative to the V-ex and N-ex groups. These unique features of W-ex mutations suggest that this mutation expression group is unlike traditional passenger or driver mutations. Based on our results, one possible explanation of why only the wild type allele is expressed in genes containing a W-ex mutation is that this particular mutation impairs the functionality or translation of a gene that is important to the tumor’s progression. Thus, the tumor selectively silences the mutant allele and only expresses the wild type allele to maintain its phenotype. A potential example of this phenomena is the mutated gene LDHB in sample 810CL. A recent study that performed an RNA interference screen in KRAS dependent lung adenocarcinomas identified that LDHB was a strong regulator of cell proliferation in these tumors^52^. LDHB has also been shown to be responsible for altering the metabolic addiction in PDAC^53^. Interestingly, the mutated LDHB gene in sample 810CL had a DNA allele frequency of 62% and 1,587 reads that aligned to the wild type allele but 0 reads that aligned to the mutant allele. This supports the evidence that the non-mutated LDHB gene is potentially a critical aspect within tumor sample 810CL. Additional confidently classified W-ex mutations from the cell line data that were noted to act in a similar manner as LDHB and have been shown to positively influence tumor progression are BCLAF1, JAK2, and KMT2D^54-56^. However, to identify exactly what impact W-ex mutations have on tumor progression, additional research is necessary. In regards to how the mutant allele of W-ex mutations is repressed, we attempted to identify if this event was due to exon skipping or nonsense mediated decay due to having a higher proportion of nonsense and indel mutations^57^. Interestingly, neither of these mechanisms seemed to be involved in the silencing of the mutant allele. Based on our statistical testing between the first exon methylation of V-ex mutations and W-ex mutations, we would suggest that the most probable mechanism of W-ex mutant allele repression is hemimethylation^46^. However, detailed experiments focused on specific loci will be important to confirm this speculation.

By classifying PDAC mutations based on their allelic expression, we were able to identify approximately 50% of all mutations within this study as likely passenger mutations, increase the sensitivity of mutation detection methods, significantly enhance the prediction of PDAC prognosis using mutation data, and identify subtypes that are statistically sensitive to specific drug therapeutics in cell culture models. These results support the concept that both V-ex and W-ex mutations contribute to the biological features of the tumor, while N-ex mutations have little influence on the progression of the tumor. Thus, we conclude that the underutilized technique of allelic expression analysis of tumor mutations provides an effective top-level method to separate mutations based on their respective tumor impact. Based on these results within PDAC, we expect that allele expression analysis will be able to assist in other aspects of cancer biology. For example, this approach can be used to explore cancer epitope detection^58^ and interrogation of tumor clonality^58,59^. Because the MAXX pipeline enables accurate identification of allelic expression for all mutation types by increasing RNA-sequencing alignment, it represents the most inclusive method to perform mutation allelic expression analyses. Custom gene specific reference genomes generated via MAXX will become a prevalent aspect in performing mutation allelic expression analyses and could be widely employed to classify mutations for advancement of precision medicine.

## Methods

### Expanded overview of MAXX pipeline

MAXX is a freely available tool (https://github.com/Adglink/MAXX) that generates gene specific reference genomes. As input, MAXX requires a mutation file, a reference genome, and the reference genome’s corresponding GTF file. The reference files used in this study were GENCODE v19 for the patient derived models and GENCODE GRCh v21 for the TCGA data^60^. In brief, MAXX uses information from the GTF file to identify the sequence of the input genes within the fasta file. Once the sequence of the genes within the mutation list is identified, MAXX first uses the gene name to create a header tag (e.g. >KRAS). Then the header tag and wild type sequence are written to the new fasta file. Next, the mutated sequence and its header are generated. The mutation sequence is created based on the information from the mutation file while the header is the gene’s name with a _Mut tag (e.g. >KRAS_Mut). The _Mut tag allows reads to be separated based on their alignment to either the wild type or mutated sequence. In cases where a gene contains multiple mutations, all mutations are inserted into the gene via a shifting algorithm. Because this study emphasized the allelic expression of mutations, MAXX is designed to only output the wild type and mutant sequence of genes presented in the mutation file. This approach dramatically decreases the size of the fasta file, which allows storage space and alignment run time to be more efficient compared to the Hg19 reference genome (Supplementary Fig. 1), while maintaining accurate RNA-sequencing alignment (Fig. 2c). In addition to the custom-made reference genome, MAXX also outputs an index file that identifies the new positions of the mutations in both the wild type and mutant sequence.

Because all of the PDAC samples within this study had undergone exome sequencing, we were able to use MAXX to generate a tumor specific reference genome for each sample. Once the new reference genome was created, Bowtie2 v. 2.3.2^61^ was used to index each reference genome. Then Tophat2 v. 2.1.1^26^ was used to align the sample’s corresponding RNA-sequencing reads. In addition to the default parameters for Tophat2, the commands “-p 10,--no-coverage-search,--b2-very-sensitive” were used. To ensure the accuracy of calculated RNA-allele frequencies, the BAM files generated by Tophat2 underwent multiple filtering steps^23,32^. First, the PCR duplicates were removed, using the Picard Tools (http://broadinstitute.github.io/picard/) MarkDuplicates command where the input parameter REMOVE_DUPLICATES was set to “true”. Second, SAMtools v. 1.4^62^ was used to keep the uniquely aligned reads via the command “samtools view-q 50-b input.bam > output.bam”. Finally, because our data was paired-end, the additional step of singleton removal using the SAMtools v. 1.4 command “samtools view-F 8-b input.bam > output.bam” was performed. The filtered bam file, tumor specific reference genome, and tumor mutation index file were then used as input for Bam-Readcount (https://github.com/genome/bam-readcount), which identified the number of reads that aligned to the mutant and wild type sequences.

### RNA mutant allele frequency calculation

To determine RNA mutant allele frequencies from the Bam-Readcount output, the number of reads that aligned to the mutant position of the mutated sequence was first calculated, then divided by the sum of the reads that aligned to the mutant position of both the wild type and mutant sequences. Due to the shifting of nucleotides from indel mutations, there were slight variations in the way the reads were calculated for the wild type allele and mutant allele between the different mutation types. For SNV mutations, the same position for the mutant allele and wild type allele was used to calculate the number of reads for each allele. As for deletion mutations, the wild type allele reads were calculated based on the average number of reads mapped at each deleted nucleotide, while the first position of the non-deleted nucleotide was used to calculate the number of mutant allele reads. To calculate the wild type and mutant allele read count for insertions, the average read count of reads aligned to each inserted nucleotide was used for the mutant allele, while the first position of the non-inserted nucleotide was used to calculate the number of reads for the wild type allele.

### Mutant expression classification expounded

Based on the variant’s total read count and RNA mutant allele frequency, the mutation was placed into a mutation expression group (Fig. 1b). To prevent misalignments and poor sequencing depth at specific regions from biasing mutation expression group placement, mutations that had a total number of reads between 3 and 9 were discarded from the study. It was also noted that a cut off of the number of reads that aligned to the mutant allele was required to differentiate W-ex mutations from V-ex mutations. Rather than have a distinct read count cutoff, the RNA mutant allele frequency of 2.5% was used to distinguish V-ex mutations from W-ex mutations.

### Patient-derived cell lines and PDX models

The establishment and variant calls of patient-derived cell lines and PDX models employed have been previously described^24,63^. The samples were exome sequenced and variant calls were identified relative to the normal tissue from the patient from which the models were derived. There is a high level of conservation between the mutations present in the models and that shown in the primary tumor.

### Sequencing of samples

The patient-derived cell lines and PDX models were subjected to exome sequencing and RNA sequencing as previously published^24,63^.

### Visualization of exome and RNA-sequencing reads

Bam files derived from exome or RNA sequencing alignment were uploaded into the Integrative Genomic Viewer software v. 2.3.93^64^. The parameters filter duplicate reads and secondary alignments were set to true.

### Generation and clustering of networks

The Cytoscape^40^ environment, in conjunction with the Reactome database, was used to generate the networks presented in this study. After downloading the ReactomeFI^41^ plugin for Cytoscape, our gene lists were uploaded using the “Gene Set/Mutation Analysis” option. The 2016 Reactome FI network version was used to visualize the results. The clustering analysis performed on the subtype networks was accomplished using the ReactomeFI plugin “Cluster FI Network” option. Only the clusters that contained at least 10% of the genes within the network were analyzed. GO Terms and associated p-values for all networks were calculated via the ReactomeFI plugin “Analyze Module Functions” option. Network visualization was enhanced using the R package ggnet2.

### Driver detection methods

As stated previously, the driver detection methods 2020+^19^, Muffin^17^, and OncodriveFM^15^ were selectively chosen because of their capability to handle indel mutations and relatively small sample sizes. Each method was run two times using either the 19 patient derived cell line data or a combination of the 19 patient derived cell line data and 75 samples from TCGA. In addition to missense, frame shift del, frame shift ins, and nonsense mutations, silence mutations were included into the input data for 2020+ and OncodriveFM, as recommend. 2020+ was ran using the cancer type specific analysis procedure found at http://2020+.readthedocs.io. Muffin was ran using the default parameters and the results produce by NDMAX with the Humannet network were used. Finally, OncodriveFM use ran with the parameter “--gt 1” and as input the functional impact scores produced by SIFT^14^, PolyPhen-2^13^, Vest^65^ and Chasm^66^ were used.

### MAXX mutation expression profiles for TCGA data

To generate a tumor specific reference genome for each sample within the TCGA-PAAD project, the somatic mutation data was obtained by downloading the PDAC Mutect2 annotation file from the website https://portal.gdc.cancer.gov/repository. This file, containing a list of all mutations for each TCGA PDAC sample, was used to generate MAXX appropriate mutation files for all available samples. The mutation files, GRCh38 v21 reference genome and GTF file were then used to generate tumor specific genomes via MAXX. Because the TCGA RNA-sequencing data needed to be re-aligned to the tumor specific genome, the RNA-seq fastq files were downloaded from https://portal.gdc.cancer.gov/legacy-archive/ and aligned using Tophat2. The rest of the analysis follows the same protocol as the cell line and PDX data to calculate mutation expression groups. Due to the confounding feature of tumor purity, we focused on the 75 TCGA PDAC cases that had the highest tumor purity score and at least 10% of the mutations were classified as V-ex mutations. To assess the tumor purity of each TCGA PDAC sample, purity scores established by Yoshihara *et al.*^44^were.downloadedfromhttp://bioinformatics.mdanderson.org/main/ESTIMATE.

### Methylation analysis expounded

The IIllumina Infinium HumanMethylation450 platform level 3 generated DNA methylation files for the TCGA samples used in this study were downloaded from https://portal.gdc.cancer.gov/repository. Calculation of the first exon methylation was performed using a custom python script. The Hg19 reference genome and corresponding GTF file was first used to identify the first exon position of all genes containing a mutation; then the first exon methylation percentage was determined by taking the average beta values for all CpG nucleotides within the first exon. Genes with mutations that were identified to be within multiple mutation expression groups between TCGA samples were excluded from the methylation analysis, as well as mutations that were classified as “discarded” (Fig 1b). Also, it was observed that approximately half of the genes within each mutation expression group did not contain first exon methylation data. This is expected to be because the CpG methylation was not available or analyzed in the first exon of these genes within the downloaded data.

### Network based stratification analysis

The NBS software was used to stratify the cell line and TCGA samples based on mutations associated with each combination of mutation expression groups. As previously mentioned, the NBS algorithm was developed by Hofree *et al.*^47^ but the implementation of NBS by Morvan *et al.*^48^ was used to stratify our samples. The source code for this NBS implementation can be found at the github repository https://github.com/marineLM/NetNorM. Multiple methods of stratification are provided, but to replicate the NBS method, the *stratification_NMF.py* script was used. This script requires a sample mutation matrix and a node edge matrix. The PathwayCommonsv6network^67^(Commons.6.All.EXTENDED_BINARY_SIF.tsv) was used to generate the node edge matrix, while different combinations of mutation expression groups were used to generate the sample mutation matrix. To replicate our results, the following commands should be used as input for the s*tratification_NMF.py* function: method_rep = ‘smoothing’, method_norm = ‘qn’, k = ‘NA’, alpha = ‘.5’, randomized = ‘False’, rs_rand = ‘NA’, and N = ‘3’.

### Survival curves

Survival information for the TCGA PDAC samples was obtained from the nationwideshildrens.org_clinical_patient_paad.txt file, downloaded from https://portal.gdc.cancer.gov/legacy-archive/ on September 6, 2017. The R package survival was then used to calculate the log rank statistic, log rank p-value, and generate the Kaplan Meier plot

### Data Availability

The MAXX software, which generates specific references genomes based on a mutation list is available for download at https://github.com/Adglink/MAXX. All RNA and exome sequencing fastq files will be available on GEO before publication of the paper.

## Acknowledgements

The authors thank all members of the Knudsen and Witkiewicz laboratory for thought-provoking discussion, technical assistance and help with manuscript preparation. This project was supported by the National Institute of Health grant R01CA211878.

## Author Contributions

ADG is responsible for creating the MAXX pipeline, data generation and writing most of the manuscript under the mentorship of ESK and AKW. ESK, AKW, and MP greatly assisted in the development of experiments which were performed in this study. PV provided fundamental scripts to perform gene expression and survival analyses. PV also contributed to the development and edits of the study.

